# State Change Probability: A Measure of the Complexity of Cardiac RR Interval Time Series Using Physiological State Change with Statistical Hypothesis Testing

**DOI:** 10.1101/817650

**Authors:** Hsuan-Hao Chao, Han-Ping Huang, Sung-Yang Wei, Chang Francis Hsu, Long Hsu, Sien Chi

## Abstract

The complexity of biological signals has been proposed to reflect the adaptability of a given biological system to different environments. Two measures of complexity—multiscale entropy (MSE) and entropy of entropy (EoE)—have been proposed, to evaluate the complexity of heart rate signals from different perspectives. The MSE evaluates the information content of a long time series across multiple temporal scales, while the EoE characterizes variation in amount of information, which is interpreted as the “state changing,” of segments in a time series. However, both are problematic when analyzing white noise and are sensitive to data size. Therefore, based on the concept of “state changing,” we propose state change probability (SCP) as a measure of complexity. SCP utilizes a statistical hypothesis test to determine the physiological state changes between two consecutive segments in heart rate signals. The SCP value is defined as the ratio of the number of state changes to total number of consecutive segment pairs. Two common statistical tests, the t-test and Wilcoxon rank–sum test, were separately used in the SCP algorithm for comparison, yielding similar results. The SCP method is capable of reasonably evaluating the complexity of white noise and other signals, including 1/f noise, periodic signals, and heart rate signals, from healthy subjects, as well as subjects with congestive heart failure or atrial fibrillation. The SCP method is also insensitive to data size. A universal SCP threshold value can be applied, to differentiate between healthy and pathological subjects for data sizes ranging from 100 to 10,000 points. The SCP algorithm is slightly better than the EoE method when differentiating between subjects, and is superior to the MSE method.

## 1. Introduction

Biologically complex systems are regulated by vast numbers of underlying components, which interact with each other nonlinearly across multiple temporal scales, to adapt to an ever-changing environment. [1,2] Thus, the output of biological systems—physiological signals—commonly exhibits complex fluctuations. [2] Peng et al. used a state space representation to illustrate the concept of adaptive processing. [3] In brief, the state space representation describes physiological states of biological systems by a set of variables, such as heart rate, blood pressure, and posture. A sudden stress to a biological system requires the system to move state location, in order to adapt to the stress. A healthy system can easily move to an appropriate state location, while a pathological system has limited adaptability, and thus stays, or shifts to an inadequate state location. This illustrates that the dynamical changes of physiological signals may reflect the adaptability of biological systems. Thus, the complexity of physiological signals is proposed to reflect the adaptability of a given biological system [1,2]. Healthy systems would have relatively high adaptability, in comparison with pathological systems [2–4]. Pathological systems, with dysfunctional components, lose the ability to adapt to variable environments, and thus appear to lose their biological complexity. [3] Consequently, a measure of complexity is expected to assign high values to healthy systems and low values to pathological systems [2–7].

In cardiac systems, three types of heart rate signals are commonly used to study and examine complexity measures. They are RR interval (beat-to-beat time interval) time series, taken from subjects with normal sinus rhythm (NSR), congestive heart failure (CHF), or atrial fibrillation (AF). NSR subjects are healthy while CHF and AF subjects are pathological cases. Typically, the RR interval time series from NSR and CHF subjects are relatively irregular and steady, respectively. In contrast, the RR interval time series of AF episodes are as random as white noise [4]. Heart rate signal fluctuates even in the absence of external perturbation, and has been studied by stochastic processing models. [8–12] Increases or decreases in RR intervals have been assumed to be independent from each other. [13] Because heart rate signals have stochastic features, many probabilistic and statistic methods have been developed to characterize their complexity.

Many entropy-based measures have been evaluated by these RR interval time series, such as, Shannon entropy (ShanEn) [14–16], approximate entropy (ApprEn) [17], sample entropy (SampEn) [18], fuzzy entropy (FuzzEn) [19], and permutation entropy (PermEn) [20]. Nevertheless, most of these measures show higher values for AF than NSR RR interval time series, which is closer to a measure of irregularity than complexity, and, thus, fail to reflect their adaptability. Comparatively, the multiscale entropy (MSE) [2,4], MSE-based method [21–23], and entropy of entropy (EoE) [24] show that the RR interval time series from NSR subjects exhibit a higher complexity than those from CHF and AF subjects. The MSE analysis evaluates the information content of a time series across multiple temporal scales, [2–4] while the EoE method quantifies the variation of information within a time series. [24]

The MSE analysis computes the sample entropy of a coarse-grained time series over multiple temporal scales. Conceptually, sample entropy evaluates the difference between the number of repeating patterns of *m* length and the number of repeating patterns of *m* + 1 length [25], where *m* represents the embedding dimensions—usually 2. The coarse-grain process resamples the original time series to reconstruct a coarse-grained time series at the temporal scale *τ*. The MSE algorithm is formed of three steps. First, the time series of a heartbeat signal is divided into consecutive, non-overlapping segments of the same length. The length corresponds to a time scale factor *τ*, which is variable. Second, the mean of each segment is computed, and the means form the coarse-grained time series. Third, the sample entropy of the coarse-grained time series is evaluated, which yields the MSE value of the original signal, at a given scale of *τ*. The steps are repeated for different *τ* (usually between 1 and 20). However, the MSE algorithm requires at least 10^*m*^, and preferably at least 30^*m*^, data points for reliable MSE evaluation [26]. Thus, MSE analysis is not suited to short heart rate signals.

In 2017, Hsu et al. proposed the EoE method to evaluate the complexity of short RR interval time series from the perspective of “state changing.” They interpreted the mean of each segment in the MSE algorithm as a “representative state” of the RR intervals, and the evaluation of sample entropy as a procedure to characterize the degree of change in the representative states. [24] Building on this concept, Hsu et al. proposed Shannon entropy of each segment as a new kind of representative state, and then used Shannon entropy to characterize the changes in these representative states. Thus, the algorithm is referred to as entropy of entropy.

The EoE method has three steps, similar to MSE analysis. First, a time series is divided into consecutive, non-overlapping segments of the same length. The length corresponds to a variable time scale factor, *τ*. Second, the Shannon entropy of each segment is computed, and these Shannon entropies form a new time series. As Shannon entropy is a measure of the amount of information in random variables, [14] the new time series is comprised of segmental information amounts from the original time series. Third, the Shannon entropy of the new time series is evaluated, which yields the EoE value of the original signal. This step evaluates the variation in the segmental information amount of the original time series, and characterizes the equivalent dynamical change state. The higher the complexity of the signal, the higher the EoE value, and the more frequent the state changes. The steps are repeated for different *τ* (usually between 1 and 10). Note that Hsu et al. extended the concept of “multiscale” from the MSE analysis. Originally, the MSE algorithm calculated the mean of each segment, in order to construct a new time series representing the original signal at the scale *τ*. Here, the EoE algorithm computes the Shannon entropy of each segment, to construct a representative state sequence, representing the original signal at the scale *τ*. The EoE algorithm is proposed to measure the complexity of short heart rate signals in the range of 70–500 data points. However, EoE value varies as the data size changes.

Even though the MSE and EoE algorithms successfully evaluate the complexity of heart rate signals for multiple temporal scales, their measures of one common synthetic signal, white noise, are problematic. White noise is completely uncorrelated and is considered not complex. [2] Thus, the complexity of white noise is independent of temporal scale and is extremely low. However, MSE value decreases, and EoE value increases, as scale increases. The MSE and EoE analyses of white noise yielded greater values at small and large scales, respectively. In addition to white noise, periodic signals and 1/f noise are two synthetic signals also commonly used to examine complexity measures. Periodic signals are not complex, while 1/f noise is considered the most complex of the three.

This study aims to develop a new complexity measure, which is suited for long and short time series, is insensitive to data size, and exhibits reasonable evaluations of synthetic and physiological signals. According to the illustration of state space representation, the dynamical changes of physiological signals reflect the adaptability of biological systems. The MSE algorithm, which characterizes repeated patterns through signals, reveals that pattern similarity is of importance at various temporal scales. The EoE analysis shows that the complexity of heart rate signals can be measured by evaluating the degree of changes in state. Based on this, we predict that a change between two physiological states may be determined by comparing the similarities between representative quantities of segments, such as their means or medians. The statistical hypothesis test is the most appropriate, since the heartbeat is modulated by partial stochastic processing.

Accordingly, we propose a new method, state change probability (SCP), to measure complexity from the “state changing” perspective. Like the MSE and the EoE methods, the algorithm of the SCP consists of three steps. The first step of the SCP is the same as that of MSE and EoE. However, in the second step, the representative state of a segment is defined as the binary result (1 or 0) of a statistical hypothesis test of two consecutive segments. If the test shows a significant difference between the two consecutive segments, the binary result is denoted as 1, which indicates the presence of physiological state changes. Otherwise, the binary result is denoted as 0, which indicates an absence of physiological state changes. All representative states form a binary series, in which each representative state labels the physiological state change between two consecutive segments with a binary digit. Therefore, in the third step, the complexity of the original time series is defined as the average of the corresponding binary series. In other words, the proposed SCP method measures the probability of physiological state change within a given time scale, using a statistical hypothesis test. Note that the multiscale concept used in the EoE method is also adopted in the SCP method.

In the following sections, the proposed SCP measure is compared with single-scale measures (ShanEn, ApprEn, SampEn, FuzzEn, and PermEn) and multi-scale measures (MSE and EoE), using three types of synthetic signal (Gaussian white noise, 1/f noise, and periodic signals) and the RR interval time series (NSR, CHF, and AF), from the databases on PhysioNet [27], for data sizes ranging from 100 to 10,000.

## 2. Materials and Methods

### 2.1. Proposed Method: State Change Probability

#### 2.1.1. Statistical Hypothesis Test

In the SCP method, the representative state of each segment is defined as the binary result of a statistical hypothesis test on a given segment, compared with its neighbor, as noted above. The independent samples t-test (t-test) and the Wilcoxon rank–sum test (W-test) are two common statistical hypothesis tests, used to examine significant differences between two data sets with acceptable confidence levels (usually 95%, *p* < 0.05). Generally, the t-test is applied when the two data sets follow a normal distribution, otherwise, the W-test is applied. In practice, any two consecutive segments of data may or may not follow a normal distribution. Therefore, in this work, the t-test and the W-test are independently applied to the SCP for every RR interval time series, for comparison.

#### 2.1.2. Algorithm

The algorithm of the SCP method using the t-test consists of three steps, as illustrated in Figure 1. First, an RR interval time series is divided into consecutive, non-overlapping segments of the same length (termed the scale factor *τ*), as shown in Figure 1a. Second, every two consecutive segments are analyzed by the t-test. The binary result of this serves as the representative state of the latter segment. As shown in Figure 1b, the value of each representative state, either 0 or 1, is color-coded, using a blank bar or a blue bar to denote the respective absence or presence of physiological state changes between the pair of consecutive segments. Thus, all the representative states form a binary series. Third, the average of the binary series is taken as the complexity value of the original time series.

**Figure 1.**
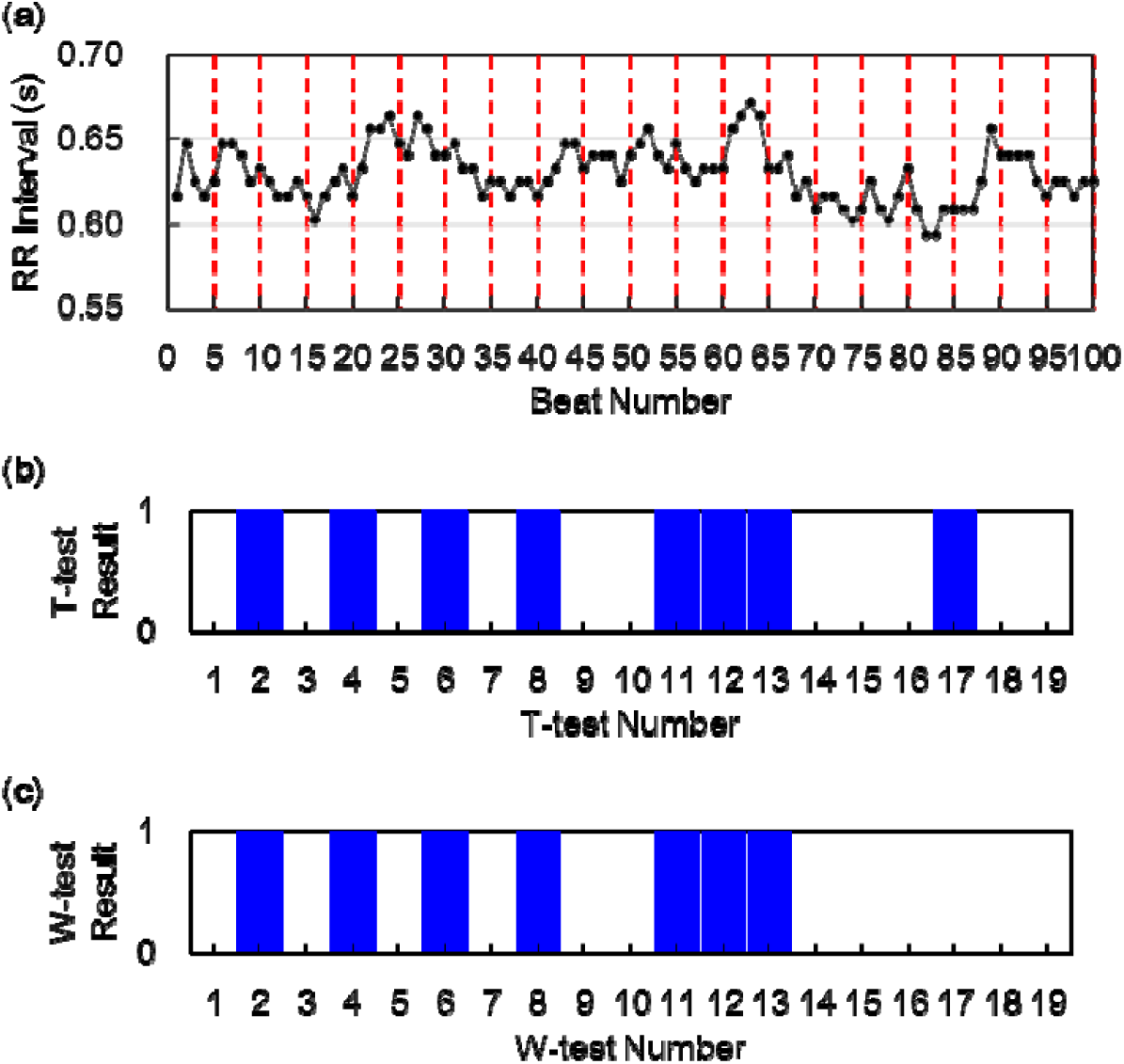
Evaluation of State Change Probability. (**a**) A cardiac RR interval time series of 100 beats. The dashed lines equally divide the time series into 20 segments. Each segment contains five beats, at a scale factor of 5. (**b**) A color-coded binary result using the t-tests on each pair of consecutive segments. (**c**) A color-coded binary result using the W-tests on each pair of consecutive segments. A blue bar represents the presence of a state change between the paired segments, while a blank bar represents the absence of a state change.

In Figure 1a, the time series consists of 100 beats, which are divided into 20 segments, at a scale factor of 5. Figure 1b illustrates the binary result of the t-test on each of the 19 pairs of consecutive segments. In this case, eight of the 19 bars are blue. Consequently, the complexity value of the original time series is 8/19 (0.42), the average of the binary series.

For comparison, the t-test is replaced by the W-test in the same SCP algorithm. Figure 1c shows the binary result of the W-test on each of the same 19 pairs. As shown in the figure, seven of the 19 bars are blue. Thus, the complexity value of the same time series using the W-test is 7/19 (0.37).

### 2.2. Data Description

The SCP method was applied to the analysis of synthetic noises and heartbeat signals. Three types of synthetic noises—Gaussian white noise (GWN), 1/f noise, and periodic signals—were used for analysis. We simulated ten Gaussian white noises and ten 1/f noises, with 10,000 points each, which were divided into equal, multiple short series of 100, 500, 1000, 5000, and 10,000 points. Thus, there were 1,000 (10 × (10,000/100)) and 200 (10 × (10,000/500)) short series, for data sizes of 100 and 500, respectively. All the short series were analyzed. Regarding periodic signals, we generated two sets of sinusoidal signals separately, with periods of 10 and 20 points. Each set had ten sinusoidal signals, with phase shifts of degrees of 0, 18, 36, 54, 72, 90, 108, 126, 144, and 162. Each sinusoidal signal was 1000 points long.

The RR interval time series were taken from the following databases on PhysioNet [27]: (1) nsrdb—MIT–BIH Normal Sinus Rhythm Database, five men aged 26 to 45, and 13 women aged 20 to 50; (2) chfdb—BIDMC Congestive Heart Failure Database, 11 men aged 22 to 71, and four women aged 54 to 63; (3) chf2db—Congestive Heart Failure RR Interval Database, 29 subjects aged 34 to 79 including eight men, two women, and 19 subjects of unknown gender; (4) ltafdb—Long Term AF Database, 84 subjects with unknown ages and genders. In the ltafdb database, the atrial fibrillation episodes were extracted according to the annotations described on PhysioNet, and were used for analysis in this study. There were only 39 atrial fibrillation episode time series longer than 10,000 heartbeats, from 39 individuals, which were used for analysis.

All the RR interval time series were filtered to remove missed beat detections and premature ventricular beats, according to the annotations on PhysioNet. Further, all time series were preprocessed, to exclude artifacts and outliers with two steps. [23,28] Firstly, RR intervals less than 0.2 s, or greater than 2.0 s, were removed. Secondly, each RR interval that differed from the mean of the 40 surrounding intervals by more than 20% was excluded. Afterward, these preprocessed RR interval time series were truncated at 10,000 points and categorized into three groups, as follows: (1) NSR group (healthy); 18 individuals from nsrdb, (2) CHF group (pathologic); 44 individuals from chfdb and chf2db, and (3) AF group (pathologic); 39 individuals from ltafdb. Each preprocessed RR interval time series was then equally divided into multiple short time series of 100, 500, 1000, 5000, and 10,000 points, for analysis and comparison.

### 2.3. Performance Metrics

The performance of the SCP method, with the optimal threshold value and scale, was evaluated using five traditional indices: accuracy, specificity, sensitivity, precision, and F_1_ score. An optimization was conducted to determine the optimal SCP threshold value and scale, leading to a maximum of *Mean F1*, which took several F_1_ scores, under various conditions, into account. An RR interval time series with an SCP value greater than this threshold was considered healthy.

Accuracy in differentiating the NSR from the CHF or the AF is defined as 100% × (*TP* + *TN*)/(*TP* + *FP* + *TN* + *FN*), denoted by *CHF Acc* or *AF Acc*, respectively. Here, *TP* is the number of the CHF or AF subjects correctly identified as the pathologic subjects, *TN* is the number of the NSR subjects correctly identified as the healthy subjects, *FP* is the number of the NSR subjects falsely identified as the pathologic subjects, and *FN* is the number of the CHF or AF subjects falsely identified as the healthy subjects. Specificity is defined as 100% × *TN*/(*FP* + *TN*) and denoted by *Spe*. Sensitivity in differentiating the NSR from the CHF or the AF is defined as 100% × *TP*/(*TP* + *FN*) and denoted by *CHF Sen* or *AF Sen*, respectively. Precision in differentiating the NSR from the CHF or the AF is defined as 100% × *TP*/(*TP* + *FP*) and denoted by *CHF Pre* or *AF Pre*, respectively.

Because the numbers of the healthy (NSR) and pathological (CHF or AF) datasets are imbalanced, overall performance is evaluated by an F_1_ score, which is used to evaluate imbalanced data. The F_1_ score for differentiating the NSR from the CHF or the AF is defined as 2 *Sen* × *Pre*/(*Sen* + *Pre*) and denoted by *CHF F*_*1*_ or *AF F*_*1*_, respectively. We defined and used *Mean F*_*1*,_ to determine the optimal scale *τ* and threshold value for the SCP analysis, to differentiate the healthy from the pathological subjects. For a given scale *τ* and threshold value, the *Mean F*_*1*_ is the average of the *CHF F*_*1*_ and the *AF F*_*1*_ for data sizes 100, 500, 1000, 5000, and 10,000.

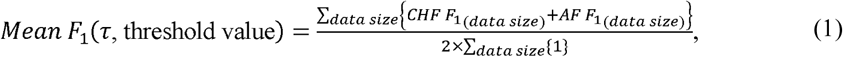

The denominator is represented as, 2 × ∑_data size_ {1} = 2 × 5 = 10, because there are five data sizes. The optimal scale and threshold value are determined when the *Mean F*_*1*_ achieves its maximum.

## 3. Results

### 3.1. Analysis of Synthetic Signals

Figure 2 shows the complexity values of four synthetic signals, evaluated by Shannon entropy (ShanEn), approximate entropy (ApprEn), sample entropy (SampEn), fuzzy entropy (FuzzEn), and permutation entropy (PermEn), respectively. The four synthetic noises are 1/f noise, Gaussian white noise (GWN), periodic signals with period of 10 data points (SinT10), and periodic signals with period of 20 data points (SinT20). As shown, the complexity values of 1/f noise (represented by a black circle) are not higher than those of GWN (represented by a blue triangle) using any of the five indices, yet the 1/f noise is considered more complex than GWN. This shows that these measures failed to accurately evaluate the complexity of the signals.

**Figure 2.**
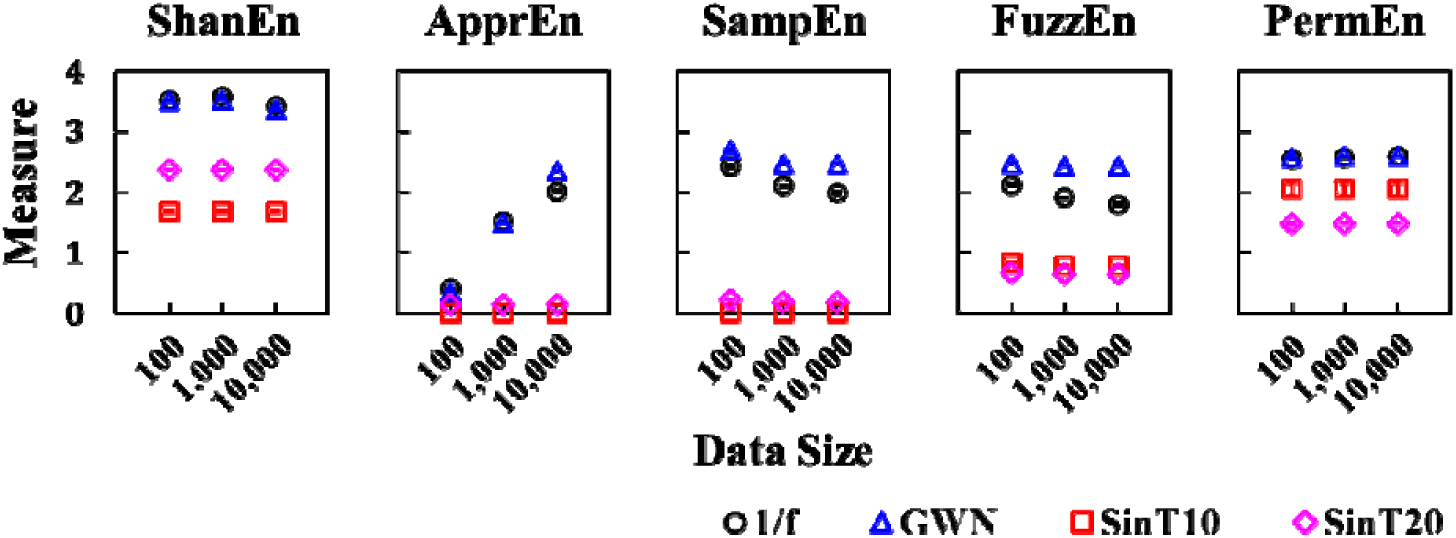
Complexity evaluations of four synthetic signals, using five single-scale measures: Shannon entropy (ShanEn), approximate entropy (ApprEn), sample entropy (SampEn), fuzzy entropy (FuzzEn), and permutation entropy (PermEn), respectively. The four synthetic signals are 1/f noise (black circle), Gaussian white noise (GWN; blue triangle), sinusoidal signals with period of 10 points (SinT10; red square), and sinusoidal signals with period of 20 points (SinT20; purple diamond). Three data sizes, 100, 1000, and 10,000, are taken into account. Error bars represent standard errors of the mean.

Figure 3 shows the complexity measures of the same four synthetic noises, evaluated using multiscale entropy (MSE), entropy of entropy (EoE), and state change probability, using the t-test (SCP_T-Test_) and W-test (SCP_W-Test_). These methods yielded the expected results, demonstrating a higher complexity value for 1/f noise, compared to white noise and periodic signals, at certain scales. For white noise, MSE and EoE analyses yielded a relatively high complexity value, while the SCP algorithm with t-test and W-test yielded a complexity value approximate to zero. The results obtained from the MSE and EoE methods are unreasonable, because white noise is completely uncorrelated and considered not complex. [2,6] Comparatively, the evaluation of the SCP algorithm is reasonable. Regarding sinusoidal signals, the MSE algorithm gives a correct result, while the SCP’s algorithm is limited, as the scale in the SCP algorithm must to be greater than the periods of the sinusoidal signals in order to evaluate complexity. EoE value oscillates with scale, which indicates that EoE cannot be applied to periodic signals. Note that the MSE analysis on data size 100 has failed to evaluate the complexity of the signals, because the MSE analysis requires data points greater than 10^*m*^-30^*m*^, where *m* = 2. [26]

**Figure 3.**
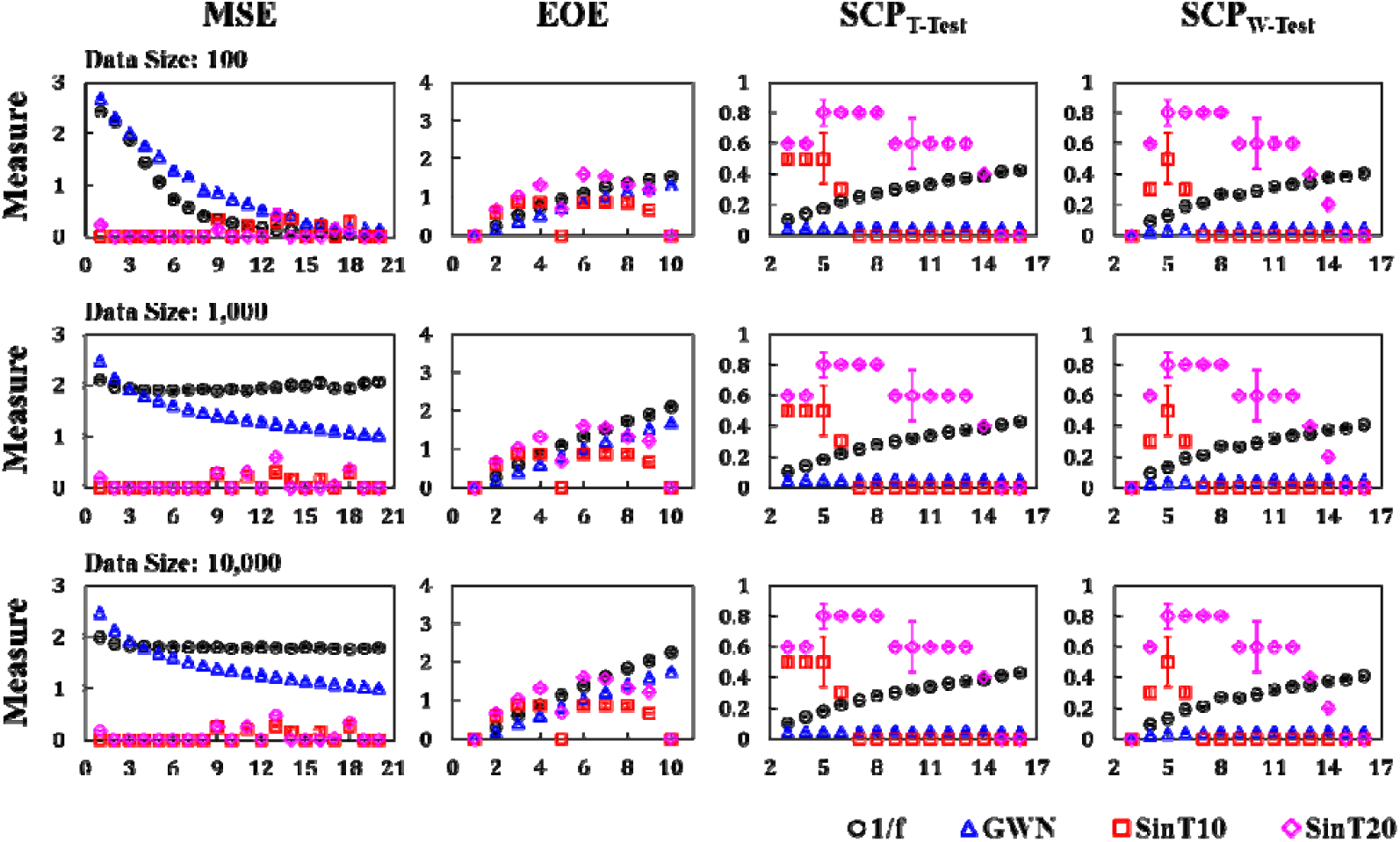
Complexity evaluations of four synthetic signals, using multiscale entropy (MSE), entropy of entropy (EoE), state change probability using the t-test (SCP_T-Test_), and state change probability using the W-test (SCP_W-Test_). The ranges of temporal scales are 1–20 for MSE, 1–10 for EoE, and 3–16 for SCP. The four synthetic signals are 1/f noise (black circle), Gaussian white noise (GWN; blue triangle), sinusoidal signals with period of 10 points (SinT10; red square), and sinusoidal signals with period of 20 points (SinT20; purple diamond). Three data sizes—100, 1000, and 10,000—are taken into account. Error bars represent standard errors of the mean.

### 3.2. Analysis of Physioloigcal Signals

Figure 4 shows the complexity measures of three types of RR interval time series, evaluated by ShanEn, ApprEn, SampEn, FuzzEn, and PermEn, respectively. The RR interval time series are taken from healthy (NSR) subjects, and subjects with atrial fibrillation (AF) or congestive heart failure (CHF). The NSR RR interval time series are considered more complex than AF and CHF time series. However, the complexity values of NSR time series obtained from these indices are not higher than those of AF and CHF. Furthermore, these indices are dependent on data size.

**Figure 4.**
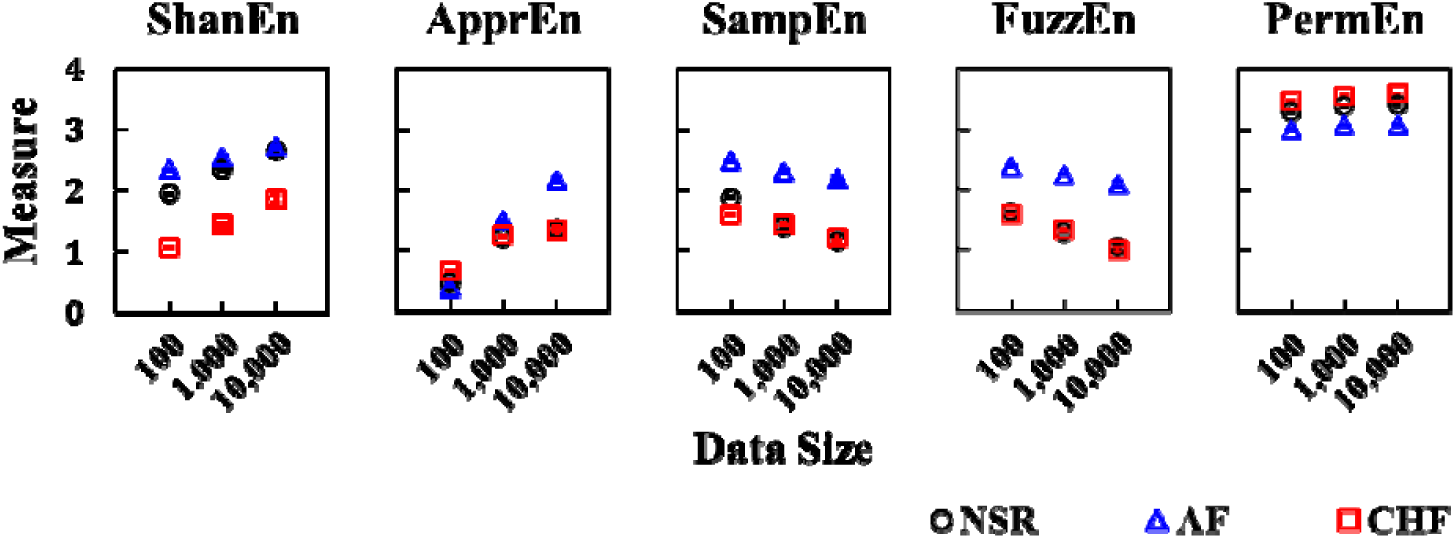
Complexity evaluations of three types of RR interval time series, using ShanEn, ApprEn, SampEn, FuzzEn, and PermEn, respectively. The RR interval time series are taken from healthy subjects (NSR; black circle), and subjects with atrial fibrillation (AF; blue triangle) or congestive heart failure (CHF; red square). Three data sizes—100, 1000, and 10,000—are taken into account. Error bars represent standard errors of the mean.

Figure 5 shows the complexity values of the same RR interval time series, evaluated by MSE, EoE, SCP_T-Test_, and SCP_W-Test_. The complexity values of the RR interval time series of healthy subjects are higher than those of the pathological subjects (AF and CHF) on some scales, which is consistent with the results of previous studies. [2,4,24] The MSE and EoE curves for the three data sizes are different from each other, while the SCP curves are insensitive to data size.

**Figure 5.**
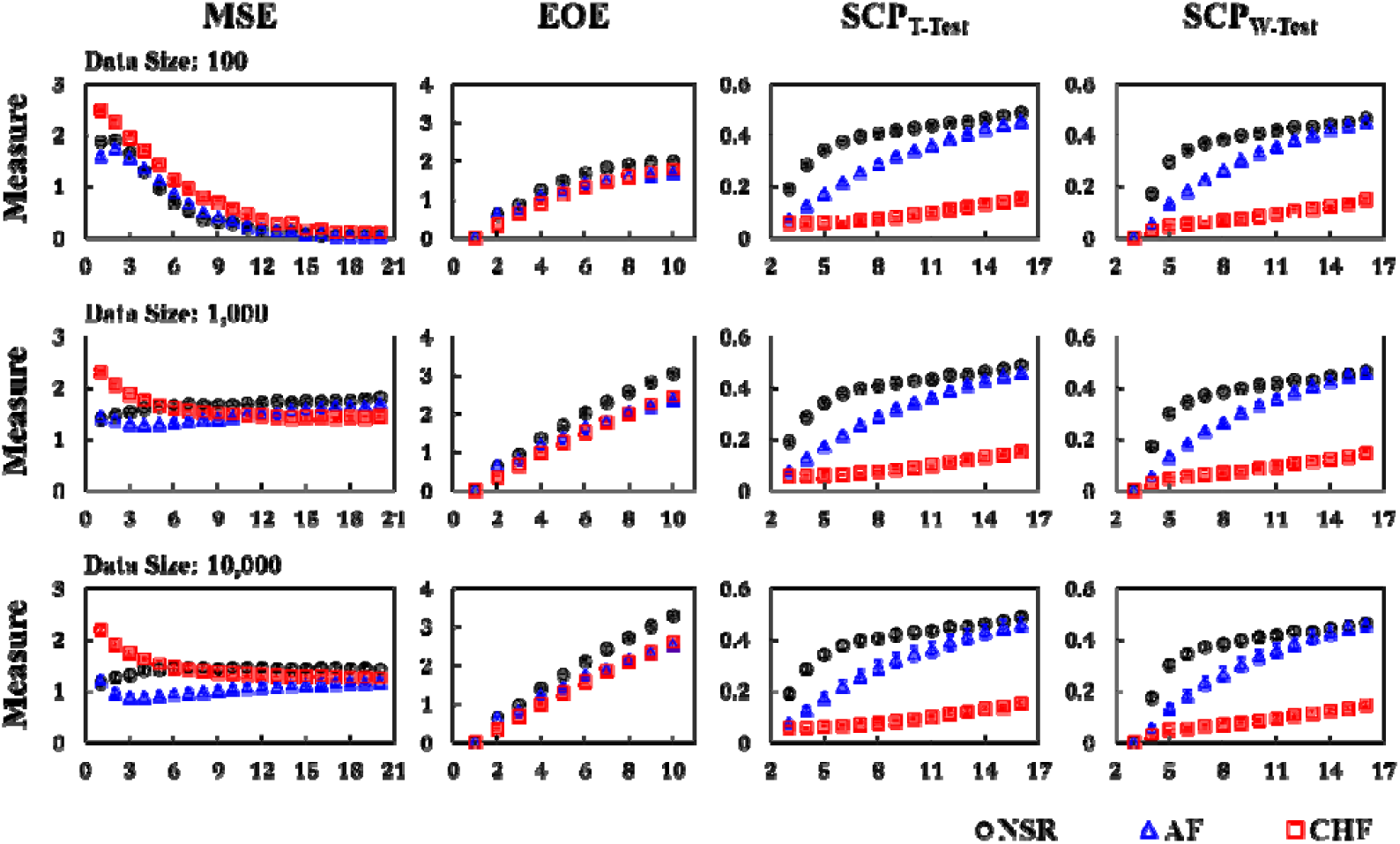
Complexity evaluations of RR interval time series, taken from NSR (black circle), AF (blue triangle), and CHF (red square) subjects, using MSE, EoE, SCP_T-Test_, and SCP_W-Test_. The ranges of scales are 1–20 for MSE, 1–10 for EoE, and 3–16 for SCP. Three data sizes—100, 1000, and 10,000—are taken into account. Error bars represent standard errors of the mean.

### 3.3. Comparison

The optimal scale and threshold value of the SCP method for analyzing the RR interval time series are determined by maximizing *Mean F*_*1*_. The optimal scale and threshold value of SCP are found to be 3 and 0.13 when using the t-test, and 4 and 0.10 when using the W-test, respectively. These optimal parameters for the SCP method are applied to different data sizes—100, 500, 1000, 5000, and 10,000—for comparison with the MSE and EoE methods. The scale used in MSE is 20 [4], and in EoE the scale is five [24].

Table 1 shows the nine performance metrics differentiating between healthy and pathological subjects. The results of ShanEn, ApprEn, SampEn, FuzzEn, and PermEn are not shown, because they failed to differentiate between healthy and pathological subjects. Their *Mean F*_*1*_ achieved a maximum of *Spe* 0%, *CHF Sen* 100%, and *AF Sen* 100%. For the same reason, the MSE analysis for data sizes 100, 500, 1000, and 5000 are not shown in Table 1. The performance metrics of the EoE and the SCP methods improve as data size increases. Note that the threshold value of EoE varies from 1.54, for data size 100, to 1.68, for data size 10,000, while the SCP threshold is universal for data sizes ranging from 100 to 10,000.

**Table 1.**
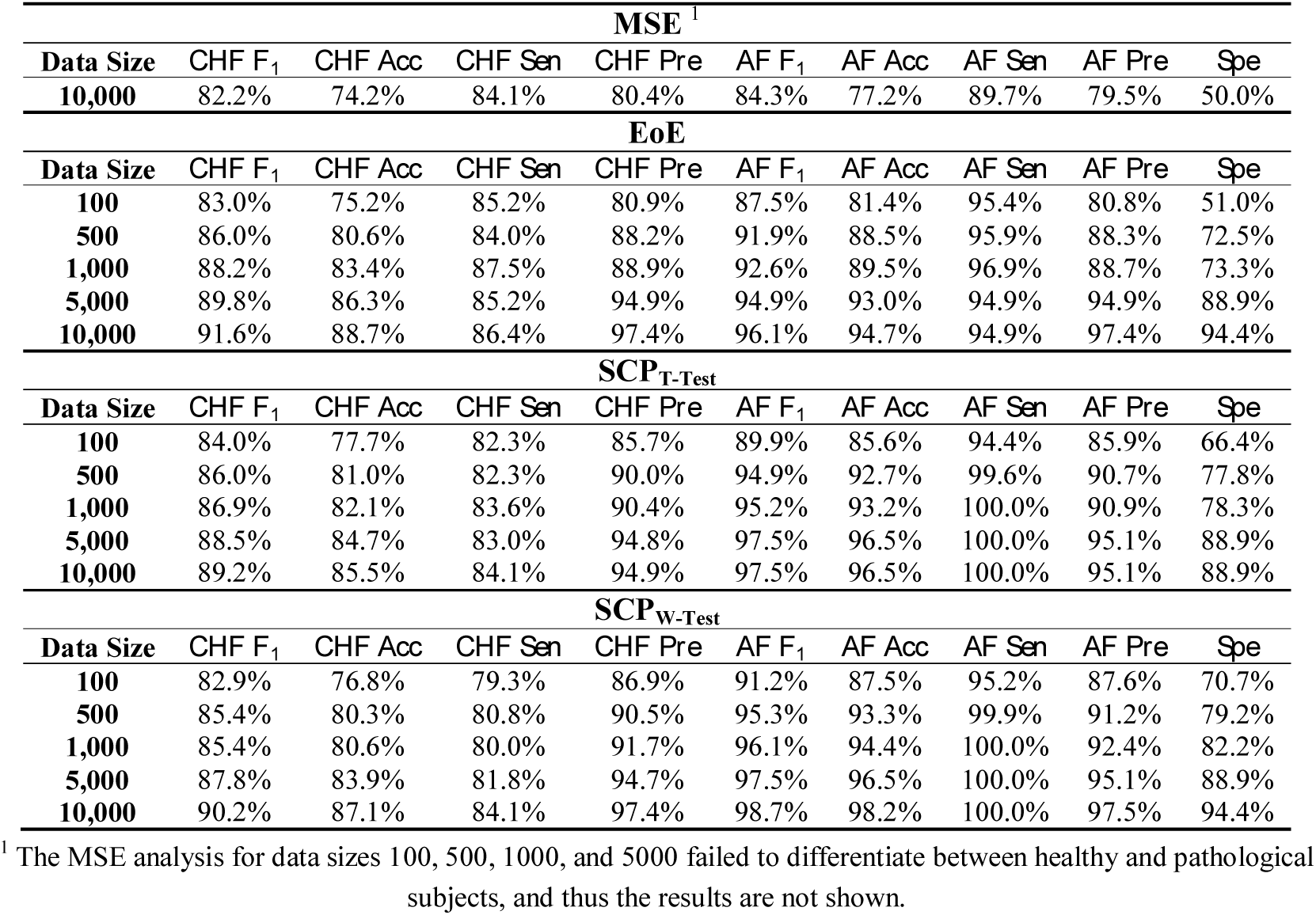
Performance when discriminating between healthy and pathological subjects using MSE, EoE, SCP_T-Test_, and SCP_W-Test_.

| **MSE<sup>1</sup>** |  |  |  |  |  |  |  |  |  |
| --- | --- | --- | --- | --- | --- | --- | --- | --- | --- |
| **Data Size** | **$CHF F_1$** | **$CHF Acc$** | **$CHF Sen$** | **$CHF Pre$** | **$AF F_1$** | **$AF Acc$** | **$AF Sen$** | **$AF Pre$** | **$Spe$** |
| **10,000** | 82.2% | 74.2% | 84.1% | 80.4% | 84.3% | 77.2% | 89.7% | 79.5% | 50.0% |
| **EoE** |  |  |  |  |  |  |  |  |  |
| **Data Size** | **$CHF F_1$** | **$CHF Acc$** | **$CHF Sen$** | **$CHF Pre$** | **$AF F_1$** | **$AF Acc$** | **$AF Sen$** | **$AF Pre$** | **$Spe$** |
| **100** | 83.0% | 75.2% | 85.2% | 80.9% | 87.5% | 81.4% | 95.4% | 80.8% | 51.0% |
| **500** | 86.0% | 80.6% | 84.0% | 88.2% | 91.9% | 88.5% | 95.9% | 88.3% | 72.5% |
| **1,000** | 88.2% | 83.4% | 87.5% | 88.9% | 92.6% | 89.5% | 96.9% | 88.7% | 73.3% |
| **5,000** | 89.8% | 86.3% | 85.2% | 94.9% | 94.9% | 93.0% | 94.9% | 94.9% | 88.9% |
| **10,000** | 91.6% | 88.7% | 86.4% | 97.4% | 96.1% | 94.7% | 94.9% | 97.4% | 94.4% |
| **$SCP_{T-Test}$** |  |  |  |  |  |  |  |  |  |
| **Data Size** | **$CHF F_1$** | **$CHF Acc$** | **$CHF Sen$** | **$CHF Pre$** | **$AF F_1$** | **$AF Acc$** | **$AF Sen$** | **$AF Pre$** | **$Spe$** |
| **100** | 84.0% | 77.7% | 82.3% | 85.7% | 89.9% | 85.6% | 94.4% | 85.9% | 66.4% |
| **500** | 86.0% | 81.0% | 82.3% | 90.0% | 94.9% | 92.7% | 99.6% | 90.7% | 77.8% |
| **1,000** | 86.9% | 82.1% | 83.6% | 90.4% | 95.2% | 93.2% | 100.0% | 90.9% | 78.3% |
| **5,000** | 88.5% | 84.7% | 83.0% | 94.8% | 97.5% | 96.5% | 100.0% | 95.1% | 88.9% |
| **10,000** | 89.2% | 85.5% | 84.1% | 94.9% | 97.5% | 96.5% | 100.0% | 95.1% | 88.9% |
| **$SCP_{W-Test}$** |  |  |  |  |  |  |  |  |  |
| **Data Size** | **$CHF F_1$** | **$CHF Acc$** | **$CHF Sen$** | **$CHF Pre$** | **$AF F_1$** | **$AF Acc$** | **$AF Sen$** | **$AF Pre$** | **$Spe$** |
| **100** | 82.9% | 76.8% | 79.3% | 86.9% | 91.2% | 87.5% | 95.2% | 87.6% | 70.7% |
| **500** | 85.4% | 80.3% | 80.8% | 90.5% | 95.3% | 93.3% | 99.9% | 91.2% | 79.2% |
| **1,000** | 85.4% | 80.6% | 80.0% | 91.7% | 96.1% | 94.4% | 100.0% | 92.4% | 82.2% |
| **5,000** | 87.8% | 83.9% | 81.8% | 94.7% | 97.5% | 96.5% | 100.0% | 95.1% | 88.9% |
| **10,000** | 90.2% | 87.1% | 84.1% | 97.4% | 98.7% | 98.2% | 100.0% | 97.5% | 94.4% |

<sup>1</sup> The MSE analysis for data sizes 100, 500, 1000, and 5000 failed to differentiate between healthy and pathological subjects, and thus the results are not shown.

For short time series, with data sizes 100 and 500, the SCP method is slightly better than EoE. For data size 100, the *CHF F*_*1*_ is 83.0% and the *AF F*_*1*_ is 87.5% for EoE; the *CHF F*_*1*_ is 84.0% and the *AF F*_*1*_ is 89.9% for the SCP using the t-test; while the *CHF F*_*1*_ is 82.9% and the *AF F*_*1*_ is 91.2% for the SCP using the W-test. For data size 500, the *CHF F*_*1*_ is 86.0% and the *AF F*_*1*_ is 91.9% for EoE; the *CHF F*_*1*_ is 86.0% and the *AF F*_*1*_ is 94.9% for the SCP using the t-test; while the *CHF F*_*1*_ is 85.4% and the *AF F*_*1*_ is 95.3% for the SCP using the W-test.

For long time series, with a data size 10,000, the results of the SCP and EoE methods are comparable, and are superior to the MSE method. The *CHF F*_*1*_ and the *AF F*_*1*_ of MSE are only around 82% and 85%, while those of SCP and EoE are around 90% and 97%, respectively.

## 4. Discussion

The SCP algorithms using the t-test and W-test led to similar results. Both successfully evaluated the complexity of signals, and were better able to differentiate between healthy and pathological subjects, compared to the MSE and EoE methods. This indicates that statistical hypothesis tests could be applied to determine changes in physiological state. Other statistical tests may also be suited for the SCP algorithm, and require further investigation.

The single-scale measures—ShanEn, ApprEn, SampEn, FuzzEn, and PermEn—failed to evaluate the complexity of synthetic and physiological signals. The 1/f noise and NSR RR interval time series are considered more complex than both white noise or periodic signals, and the CHF or AF RR interval time series. However, these indices do not yield a high value on the 1/f noise and NSR time series. By comparison, the multi-scale measures, MSE, EoE, and SCP, can reflect the complexity of these signals.

Regarding synthetic signals, MSE and EoE result in an unreasonable evaluation of Gaussian white noise. The MSE analysis of Gaussian white noise yields relatively high values at a small scale, and the MSE value decreases as the scale increases. On the contrary, the EoE analysis on Gaussian white noise yields relatively high values at larger scales, and the EoE value increases as the scale increases. Both results are incorrect, because the Gaussian white noise is completely uncorrelated and is independent of temporal scale. A reasonable complexity measurement of Gaussian white noise is approximated to zero for all scales, as is shown by the SCP algorithm.

Most of the indices are sensitive to data size, which could lead to inconvenience and difficulties in future clinical applications. The SCP algorithm is insensitive to data size. This allows for comparison between the results of SCP analyses of different data sizes.

## 5. Conclusions

From the “state changing” perspective, we propose the SCP method as a new measure of complexity in synthetic and heartbeat signals. The SCP algorithm determines state changes by means of a statistical hypothesis test. In this study, two common statistical hypothesis tests, the t-test and the Wilcoxon rank–sum test, are separately used in the SCP algorithm, yielding similar results. The use of either test showed higher SCP values for NSR signals, compared with those of CHF and AF signals, for scales in the range 4–14. Therefore, the proposed SCP method, using either the t-test or W-test, shows potential as a means of analyzing short heart rate signals.

The SCP method has three advantages compared to other measures. First, SCP analysis exhibits reasonable evaluations of the complexity of synthetic and physiological signals. Both the MSE and EoE methods result in unreasonable evaluations of white noise. When evaluating periodic signals, the SCP algorithm requires that the temporal scale *τ* is greater than the period of the signals for meaningful evaluation, while the EoE method is not suited for analyzing periodic signals. Second, the SCP algorithm can be applied to both short and long signals. When analyzing heart rate signals, 100 heart beats are sufficient for SCP to achieve at least an 82% F_1_ score when differentiating between healthy and pathological subjects, which is comparable with the EoE method and superior to the MSE analysis. Third, the SCP algorithm is insensitive to data size. Many measures, such as ShanEn, ApprEn, SampEn, FuzzEn, MSE, and EoE, are sensitive to data size. Thus, the threshold value for differentiating between the healthy and the pathological subjects of EoE shifts as the data size changes. For the SCP algorithm, the threshold values of 0.13 using the t-test, and 0.10 using the W-test, can be universally applied to data sizes ranging from 100 to 10,000 points. This is convenient for instances in which data size changes under different conditions.

## Author Contributions

Conceptualization, H.-H.C.; methodology, H.-H.C. and H.-P.H.; validation S.-Y.W. and C.F.H.; formal analysis, H.-H.C. and H.-P.H.; writing—original draft preparation, H.-H.C.; writing—review and editing, L.H. and S.C.; supervision, L.H. and S.C.

## Funding

This research was funded by the Ministry of Science and Technology of the Republic of China, grant number MOST107-2221-E009-016.

## Conflicts of Interest

The authors declare no conflict of interest.

## References

1. Mitchell, M. Complexity: A guided tour. New York: Oxford University Press, Inc. 2009.

2. Costa, M.; Goldberger, A.L.; Peng, C.K. Multiscale entropy analysis of biological signals. Phys Rev E 2005, 71.

3. Peng, C.K.; Costa, M.; Goldberger, A.L. Adaptive data analysis of complex fluctuations in physiologic time series. Adv Adapt Data Anal 2009, 1, 61–70.

4. Costa, M.; Goldberger, A.L.; Peng, C.K. Multiscale entropy analysis of complex physiologic time series. Phys Rev Lett 2002, 89.

5. Goldberger, A.L.; Amaral, L.A.; Hausdorff, J.M.; Ivanov, P.; Peng, C.K.; Stanley, H.E. Fractal dynamics in physiology: Alterations with disease and aging. Proc Natl Acad Sci U S A 2002, 99 Suppl 1, 2466–2472.

6. Goldberger, A.L.; Peng, C.K.; Lipsitz, L.A. What is physiologic complexity and how does it change with aging and disease? Neurobiol Aging 2002, 23, 23–26.

7. Lipsitz, L.A.; Goldberger, A.L. Loss of ‘complexity’ and aging. Potential applications of fractals and chaos theory to senescence. JAMA 1992, 267, 1806–1809.

8. Amaral, L.A.N.; Goldberger, A.L.; Ivanov, P.C.; Stanley, H.E. Modeling heart rate variability by stochastic feedback. Comput Phys Commun 1999, 121, 126–128.

9. Stanley, G.B.; Poolla, K.; Siegel, R.A. Threshold modeling of autonomic control of heart rate variability. Ieee T Bio-Med Eng 2000, 47, 1147–1153.

10. Kuusela, T.; Shepherd, T.; Hietarinta, J. Stochastic model for heart-rate fluctuations. Phys Rev E 2003, 67

11. Lu, S.; Kanters, J.; Chon, K.H. A new stochastic model to interpret heart rate variability. P Ann Int Ieee Embs 2003, 25, 2303–2306.

12. Kuusela, T. Stochastic heart-rate model can reveal pathologic cardiac dynamics. Phys Rev E 2004, 69.

13. Costa, M.; Goldberger, A.L.; Peng, C.K. Broken asymmetry of the human heartbeat: Loss of time irreversibility in aging and disease. Phys Rev Lett 2005, 95.

14. Shannon, C.E. The mathematical theory of communication (reprinted). M D Comput 1997, 14, 306–317.

15. Carrillo, A.E.; Flouris, A.D.; Herry, C.L.; Poirier, M.P.; Boulay, P.; Dervis, S.; Friesen, B.J.; Malcolm, J.; Sigal, R.J.; Seely, A.J.E., et al. Heart rate variability during high heat stress: A comparison between young and older adults with and without type 2 diabetes. Am J Physiol-Reg I 2016, 311, R669–R675.

16. Flouris, A.D.; Friesen, B.J.; Herry, C.L.; Seely, A.J.E.; Notley, S.R.; Kenny, G.P. Heart rate variability dynamics during treatment for exertional heat strain when immediate response is not possible. Exp Physiol 2019, 104, 845–854.

17. Pincus, S.M. Approximate entropy as a measure of system-complexity. P Natl Acad Sci USA 1991, 88, 2297–2301.

18. Richman, J.S.; Moorman, J.R. Physiological time-series analysis using approximate entropy and sample entropy. Am J Physiol-Heart C 2000, 278, H2039–H2049.

19. Chen, W.; Zhuang, J.; Yu, W.; Wang, Z. Measuring complexity using fuzzyen, apen, and sampen. Medical engineering & physics 2009, 31, 61–68.

20. Bandt, C.; Pompe, B. Permutation entropy: A natural complexity measure for time series. Phys Rev Lett 2002, 88.

21. Silva, L.E.V.; Cabella, B.C.T.; Neves, U.P.D.; Murta, L.O. Multiscale entropy-based methods for heart rate variability complexity analysis. Physica A 2015, 422, 143–152.

22. Humeau-Heurtier, A. The multiscale entropy algorithm and its variants: A review. Entropy-Switz 2015, 17, 3110–3123.

23. Costa, M.D.; Goldberger, A.L. Generalized multiscale entropy analysis: Application to quantifying the complex volatility of human heartbeat time series. Entropy-Switz 2015, 17, 1197–1203.

24. Hsu, C.F.; Wei, S.Y.; Huang, H.P.; Hsu, L.; Chi, S.; Peng, C.K. Entropy of entropy: Measurement of dynamical complexity for biological systems. Entropy-Switz 2017, 19.

25. Gutierrez-Tobal, G.C.; Alvarez, D.; Gomez-Pilar, J.; del Campo, F.; Hornero, R. Assessment of time and frequency domain entropies to detect sleep apnoea in heart rate variability recordings from men and women. Entropy-Switz 2015, 17, 123–141.

26. Pincus, S.M.; Goldberger, A.L. Physiological time-series analysis - what does regularity quantify. Am J Physiol 1994, 266, H1643–H1656.

27. Goldberger, A.L.; Amaral, L.A.N.; Glass, L.; Hausdorff, J.M.; Ivanov, P.C.; Mark, R.G.; Mietus, J.E.; Moody, G.B.; Peng, C.K.; Stanley, H.E. Physiobank, physiotoolkit, and physionet - components of a new research resource for complex physiologic signals. Circulation 2000, 101, E215–E220.

28. Costa, M.; Healey, J.A. Multiscale entropy analysis of complex heart rate dynamics: Discrimination of age and heart failure effects. Comput Cardiol 2003, 30, 705–708

